# Dimerization of human drebrin-like protein governs its biological activity

**DOI:** 10.1101/869925

**Authors:** Arindam Ghosh, Jörg Enderlein, Eugenia Butkevich

## Abstract

Drebrin-like protein (DBNL) is a multidomain F-actin binding protein, which also interacts with other molecules within different intracellular pathways. Here, we present quantitative measurements on size and conformation of human DBNL. Using dual focus fluorescence correlation spectroscopy, we determined the hydrodynamic radius of DBNL monomer. Native gel electrophoresis and dual color fluorescence cross-correlation spectroscopy show that both endogenous and recombinant DBNL exist as dimer under physiological conditions. We demonstrate that C-terminal truncations of DBNL downstream of the coiled-coil domain result in its oligomerization at nanomolar concentration. In contrast, the ADF-H domain alone is a monomer, which displays a concentration-dependent self-assembly. *In vivo* FLIM-FRET imaging shows that the presence of only actin-binding domains is not sufficient for DBNL to localize properly at actin filament inside the cell. In summary, our work provides a detailed insight on structure-function relationship of human drebrin-like protein.

## Introduction

Drebrin-like protein (also known as DBNL, mAbp1, HIP-55, SH3P7, DBN-1 in *Caenorhabditis elegans*) is an actin-binding protein of the ADF-H family. Its amino acid sequence consists of an N-terminal ADF-H domain, followed by a coiled-coil region, a proline-rich sequence, and a C-terminal SH3 domain. The multi-domain structure of DBNL allows it to serve as an adapter protein for connecting the actin cytoskeleton to many biomolecules enabling multiple cellular functions. Its interaction with F-actin is mediated via the two actin-binding modules: an ADF-H domain and a coiled-coil region^1–3^. Besides, DBNL interacts with dynamin 1, WASP-interacting protein WIP, Piccolo, the Cdc42 guanine nucleotide exchange factor Fgd1, FHL2, and other proteins^4–8^. DBNL is an important player that mediates endocytosis and vesicle recycling^4,9–12^. In addition, it regulates actin dynamics during formation of dorsal ruffles^5^ and podosomes^13^ as well as during sarcomere contraction^3^. A recent report indicates the role of native DBNL as a negative regulator of cancer development, while its ADF-H domain alone enhances Rho GTPase signaling and increases cancer cell invasion^8^. Atomic force microscopy of *C. elegans* drebrin-like protein revealed its globular shape^3^. The crystal structure of the ADF-H domain of *Saccharomyces cerevisiae* DBNL has been resolved^2^. Recently, a dimeric structure of *S. cerevisiae* DBNL bound to the Arp2/3 complex has been found using electron microscopy^14^.

In this report, we determine the size of human DBNL monomers, examine their self-organization under physiological conditions, and elucidate the impact of coiled-coil domain in this process. In addition, we demonstrate the importance of native conformation for the proper interaction of DBNL with actin filaments in cells. For this, we utilize advanced variants of fluorescence correlation spectroscopy (FCS), native gel electrophoresis, and fluorescence lifetime imaging – Förster resonance energy transfer (FLIM-FRET).

## Results and Discussion

### DBNL exists as a dimer under physiological conditions

To examine the conformational state of endogenous human DBNL, we performed native gel electrophoresis of cell lysates from four different human cell lines: HeLa (cervix epithelial adenocarcinoma), MCF7 (mammary gland epithelial adenocarcinoma), HEK-293 (embryonic kidney) and hMSC (human bone marrow derived mesenchymal stem cells). While the predicted molecular weight of human DBNL is ∼ 48 kDa, it migrates as a single band of ∼ 60 kDa under denatured conditions. Under native conditions, it migrates as a single band of ∼ 120 kDa, that strongly indicates its dimerization (Fig. 1A).

**Figure 1:**
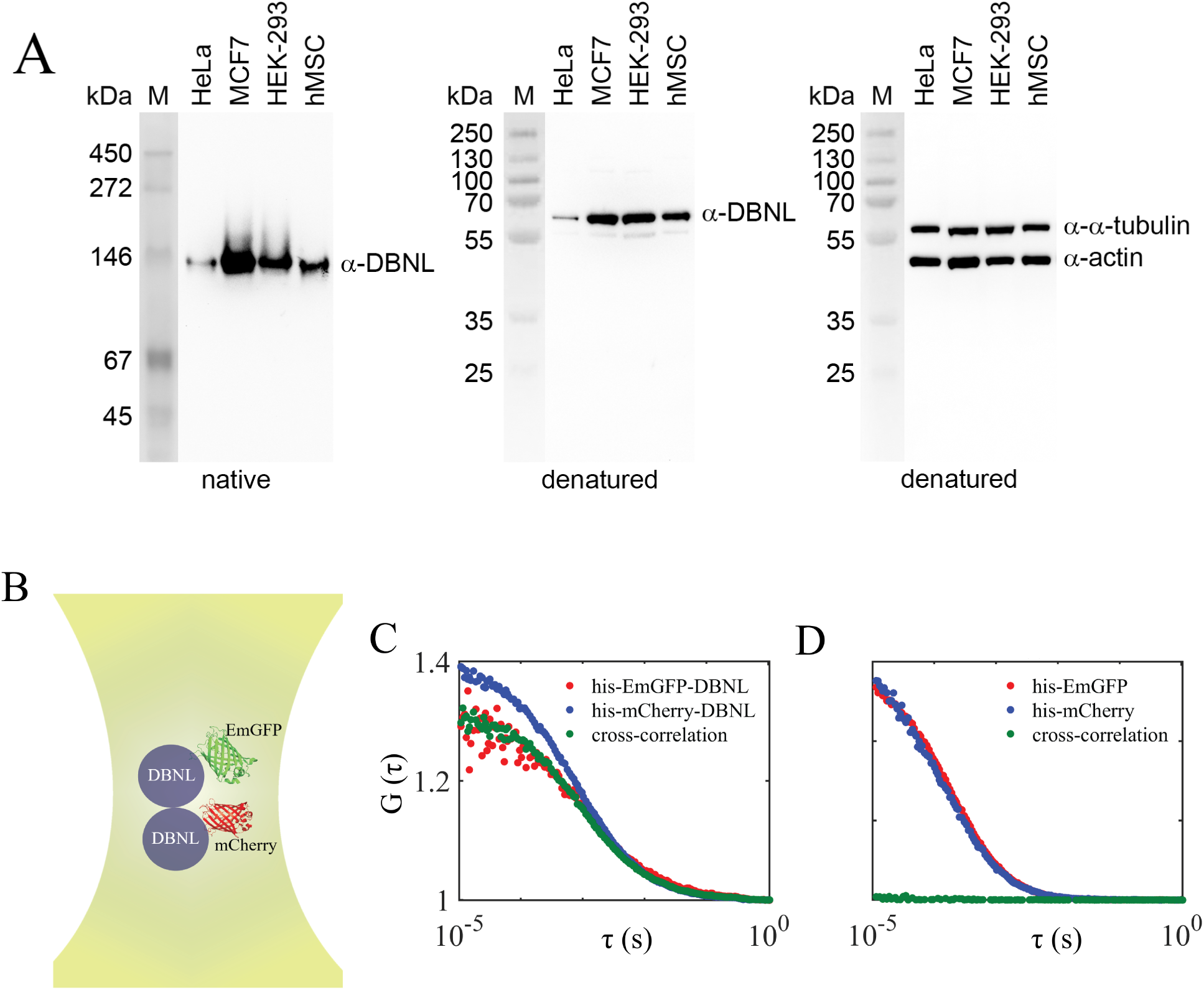
Both endogenous and recombinant DBNL occur as dimers. **A**. DBNL exists as a dimer in human cells. Western blot analysis of DBNL expression in HeLa (cervix epithelial adenocarcinoma) cells, MCF7 (mammary gland epithelial adenocarcinoma) cells, HEK-293 (embryonic kidney) and hMSC (human bone marrow derived mesenchymal stem cells). Proteins of total cell lysates were separated by native 10% Tris-Glycin or denatured 12% Bis-Tris gel electrophoresis, transferred onto a PVDF membranes and stained with anti-human DBNL antibodies or anti-*α*-tubulin and anti-actin antibody together for loading control. Molecular mass markers in kDa are indicated on the left. **B – D**. Recombinant DBNL exists as a dimer. **B**. Experimental scheme of dual-colour fluorescence cross-correlation spectroscopy (FCCS) of equimolar mixture of his-EmGFP-DBNL and his-mCherry-DBNL. Image shows an illustration of co-diffusion of DBNL monomers tagged with his-EmGFP and his-mCherry in a focused confocal volume. **C**. Autocorrelation curves of his-EmGFP-DBNL (red), his-mCherry-DBNL (blue), and their cross-correlation (green). **D**. The same for his-EmGFP and his-mCherry. As can be seen in **D**, we observe negligible or no cross-correlation between his-EmGFP and his-mCherry unlike when they are tagged to DBNL as in **C**. All correlation curves were normalized to their amplitudes at time *τ* = 1 s when the correlations have decayed completely.

Next, we investigated the dimerization of recombinant DBNL using dual colour fluorescence cross-correlation spectroscopy (FCCS). FCCS is an extension of FCS (fluorescence correlation spectroscopy) where one detects the fluorescence signal from two spectrally distinct fluorescent species simultaneously in two channels (please see Fig.1B)^15,16^. A subsequent cross-correlation analysis allows the detection of particles carrying both fluorescent labels. The method has been widely exploited to study protein-protein interactions in signaling processes, kinetics of enzymatic cleavage, or dynamic co-localization of proteins in endocytic pathways and intracellular trafficking ^17–21^. We perform FCCS measurements on an equimolar mixture of his-EmGFP-DBNL (DBNL monomer fused with his-tagged emerald green fluorescent protein) and his-mCherry-DBNL (DBNL monomer fused with his-tagged mcherry) monomers. If EmGFP- and mCherry-labeled monomeric units of DBNL bind to each other, this results in a non-zero fluorescence cross-correlation. Figure 1C shows the fluorescence auto- and cross-correlation curves obtained from a nanomolar mixture of his-EmGFP-DBNL and his-mCherry-DBNL. We observe a positive cross-correlation amplitude due to co-diffusion of DBNL monomers. In contrast, we observe negligible or no cross-correlation in case of only his-EmGFP and his-mCherry (Fig.1D). This finding is in excellent agreement with experiments on endogenous human DBNL. These observations point out that both endogenous and recombinant human DBNL undergo dimerization under physiological conditions.

### Hydrodynamic radius of DBNL

We utilized dual-focus FCS (2fFCS) to measure the translational diffusion coefficient of his-EmGFP-DBNL monomers in solution. This value was subsequently converted to hydrodynamic radius or radius of hydration of the protein using the Stokes-Einstein relation

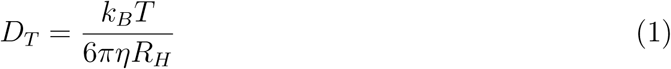

where *D*_*T*_ is the diffusion coefficient at measurement temperature *T, k*_*B*_ is Boltzmann’s constant, *η* is the viscosity of the solution and *R*_*H*_ is the Stokes or hydrodynamic radius. Thus, any change in *R*_*H*_ of a molecule of interest is directly reflected in a change of its translational diffusion coefficient. 2fFCS, in contrast to classical FCS, is not affected by refractive index mismatch, thickness variations in glass coverslip, laser beam astigmatism, or optical saturation of the fluorescent molecules. The technique was introduced in 2007 ^22^ and has been utilized for precise determination of diffusion coefficients of fluorescent molecules at pico- to nanomolar concentrations ^23–25^. Here, we briefly outline the working principle of the technique, for details, please see ^22^. A pair of interleaved pulsing lasers with orthogonal polarization are used for sample excitation. A Nomarski prism in the excitation path between the dichroic mirror and the objective lens deflects the light in a polarization-dependent manner so that after focusing through the objective two laterally shifted but overlapping excitation foci are created. The lateral distance between the foci is wavelength-dependent and has to be determined a priori by a calibration measurement with a dye or fluorescent beads with known diffusion coefficient.

This distance remains unaltered under optical saturation or aberrations, thus can be used as a “ruler” for measuring diffusion coefficients. By employing pulsed interleaved excitation (PIE) together with time-correlated single photon counting (TCSPC), one can determine which detected photon was excited by which laser and thus in which of the two laterally shifted foci. Next, we calculate the fluorescence autocorrelation function (ACF) of each focus and the cross-correlation function (CCF) between foci, and perform a diffusion coefficient global fit of the ACF and CCF using appropriate model functions^22^. For our measurements on DBNL, we used 100 pM of his-EmGFP-DBNL solution in 1x PBS (pH 7.4) and employed a commercial confocal microscope for the 2fFCS experiments. Figure 2 shows the ACFs and CCFs which were then fitted globally resulting in the following fit parameters: diffusion coefficient *D* = (37.50 ± 2.50) *µ*m^2^/s, triplet state relaxation time *τ*_*T*_ = 152.2 ± 3.5 *µ*s, laser-focus beam waist diameter *ω*_0_= 435 nm and and Rayleigh length *a*_0_ = 203 nm. Using equation 1, we calculated a Stokes radius of his-EmGFP-DBNL of 6.72 ± 0.40 nm. Next, we quantified the diffusion coefficient of his-EmGFP only (D = 104.20 ± 8.50 *µ*m^2^/s), which yields a *R*_*H*_ value of 2.42 ± 0.20 nm that matches previously reported value^26^. Hence, a difference of 4.30 ± 0.20 nm is observed between the hydrodynamic radius values of his-EmGFP-tagged DBNL and his-EmGFP. However, the linker of 14 amino acids’ length connecting the his-EmGFP and DBNL is not taken into account in this case. Approximating a fusion protein linker length of roughly 1 nm, we can interpret the hydrodynamic radius (*R*_*H*_) of DBNL monomer as ∼ 3.30 ± 0.20 nm.

**Figure 2:**
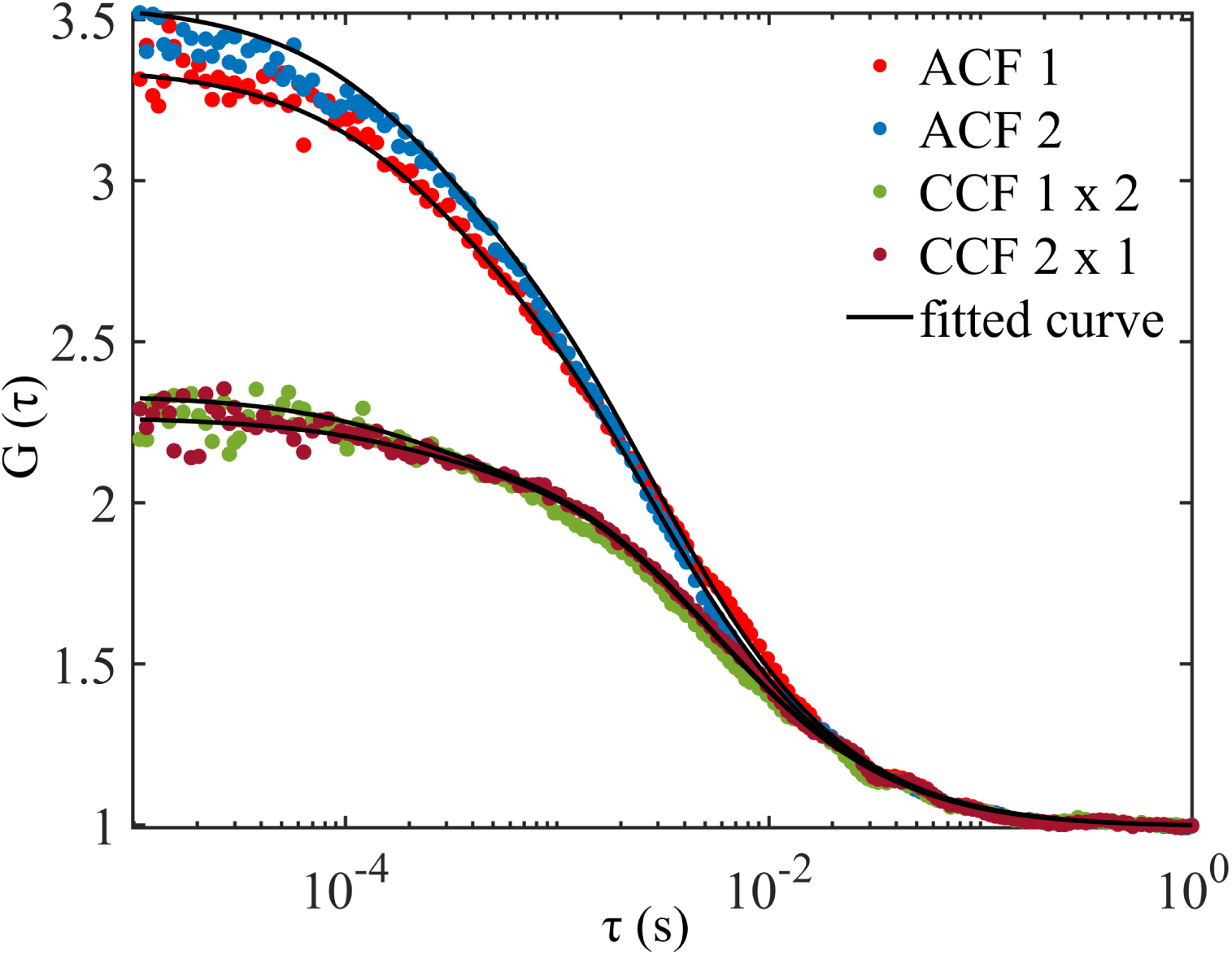
2fFCS measurement of 100 pM aqueous solution of his-EmGFP-DBNL. Autocorrelation functions (ACF) are shown as red circles for the first focus (ACF 1) and blue circles (ACF 2) for the second focus. The two possible cross-correlation functions (CCF) between both foci are represented as green and brown circles respectively. Solid lines indicate a global fit. As fit parameters, we obtained the diffusion coefficient *D* = 37.50 ± 2.50 *µ*m^2^/s, the triplet state relaxation time *τ*_*T*_ = 152.2 ±3.5 *µ*s, and *ω*_0_ = 435 nm and *a*_0_ = 203 nm. All correlation curves were normalized to their amplitudes at time *τ* = 1 s. Excitation intensity was 10 microwatts for each laser.

### Biophysical properties of truncated DBNL

The shape of a protein is specified by non-covalent interactions between regions of its amino-acid sequence. Mutations of proteins are well known to destabilize their conformation and initiate oligomerization and even aggregation, which might cause cellular dysfunction^27–29^. A two-stranded *α*-helical coiled-coil is a most common structural motif that mediates dimerization via hydrophobic and electrostatic interactions between residues^30,31^. To examine the impact of the coiled-coil domain in the dimerization of DBNL, we investigated three DBNL truncation mutants: DBNL(1-179) (truncated before coiled-coil domain), DBNL(1-256) (truncated after coiled-coil domain) and DBNL(1-374) (deleted SH3 domain), as shown in Figure 3A. The non-tagged truncated proteins were expressed in MCF7 cells under the control of a CMV promoter, and cell lysates were subjected to native gel electrophoresis and western blot analysis. Surprisingly, none of the mutated proteins displayed a monomeric or dimeric structure. Instead, they migrated through the native gel significantly slower than the full-length DBNL (Fig. 3B). Similarly, the native recombinant his-EmGFP-tagged truncated proteins showed smeared patterns with molecular weights larger than the full-length his-EmGFP-DBNL (Fig. 3C and D). These results clearly indicate that truncations of DBNL lead to a change in compaction and might induce oligomerization.

**Figure 3:**
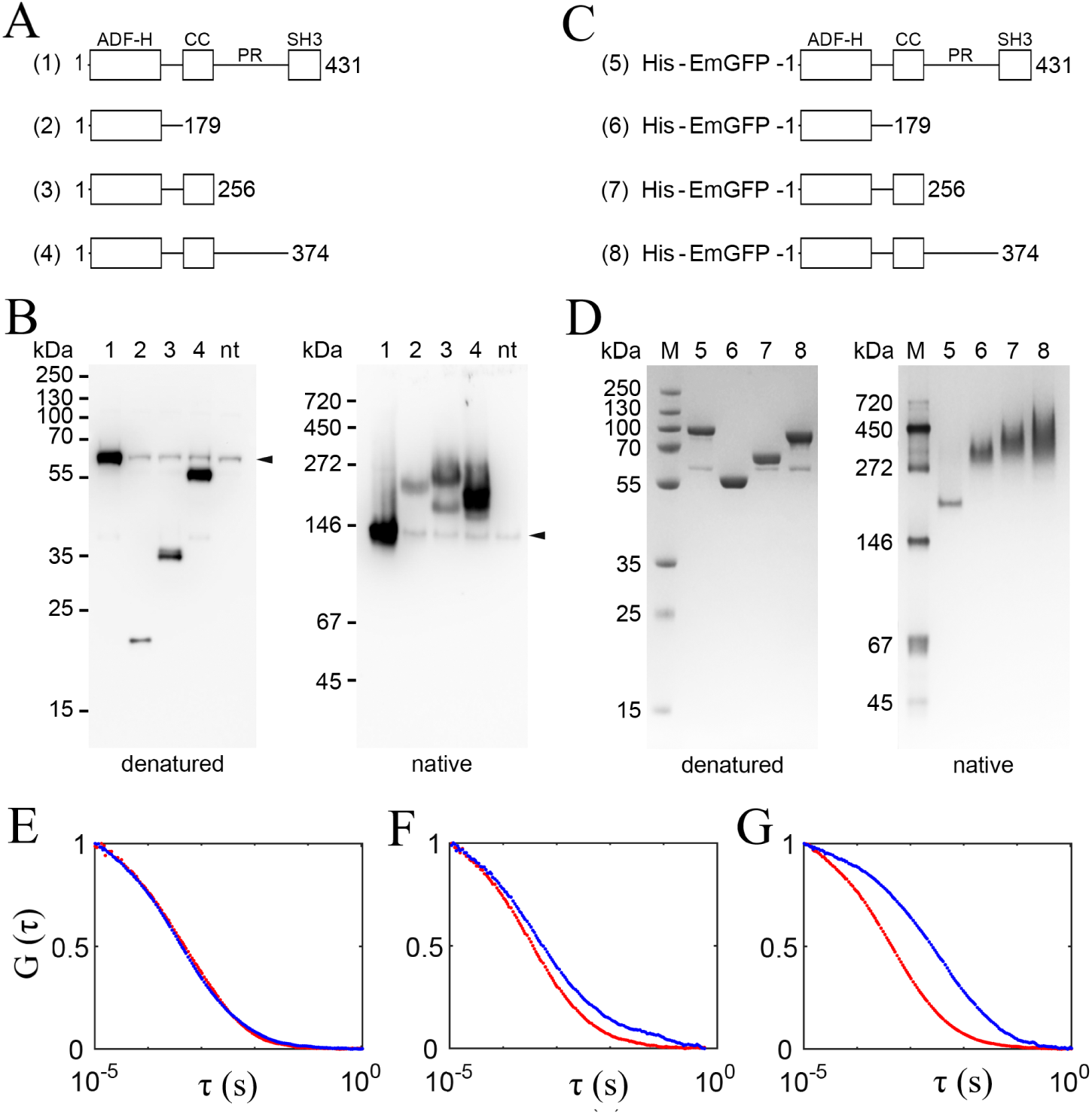
Biophysical properties of truncated DBNL. **A**. Schematic representation of full-length DBNL and its truncation mutants: DBNL(1-179), DBNL(1-256) and DBNL(1-374). **B**. Western blot analysis of DBNL in MCF7 cells transfected with full-length DBNL or its truncation mutants, indicated in **A**, and in non-transfected cells (nt). Denatured or native proteins were separated by 12% Bis-Tris or 10% Tris-Glycin gel electrophoresis respectively, transferred onto a PVDF membrane and stained with anti-human DBNL antibodies. Molecular mass markers in kDa are indicated on the left. The low-intensity bands indicated by arrowheads are of endogenous DBNL. The weak signal of DBNL(1-179) is due to the reduced number of binding sites for the polyclonal antibodies. **C**. Schematic representation of recombinant his-EmGFP-tagged full-length DBNL and its truncated mutants: DBNL(1-179), DBNL(1-256) and DBNL(1-374). **D**. Denatured or native proteins, indicated in **C**, were separated by 12% Bis-Tris or 10% Tris-Glycin gel electrophoresis respectively and stained with Roti^®^-Blue quick. Molecular mass markers in kDa are indicated on the left. **E-G**. FCS measurements on DBNL truncated at three different positions. **E** Normalized fluorescence autocorrelation of his-EmGFP-DBNL(1-179). Autocorrelation curves for 100 pM and 10 nM solutions of the protein are represented by red and blue, respectively. As can be seen, there is negligible difference in diffusion timescales for 100 pM and 10 nM. **F**. The same for his-EmGFP-DBNL(1-256). Diffusion speed slows down for 10 nM (blue) compared to 100 pM protein concentration (red). **G**. The same for his-EmGFP-DBNL(1-374). Protein oligomerization leads to slower diffusion at 10 nM (blue) relative to 100 pM (red) protein concentration.

To probe oligomerization of truncated DBNL, we applied classical FCS and measured the translational diffusion times of recombinant his-EmGFP-tagged DBNL(1-179), DBNL(1-256) and DBNL(1-374) at 100 picomolar and 10 nanomolar concentrations in 1x PBS buffer (pH 7.4) using a laser intensity of 10 *µ*W. The his-EmGFP-DBNL(1-179) showed the same diffusion times at 100 pM and 10 nM concentrations (Fig. 3E). In contrast, diffusion times of both the his-EmGFP-DBNL(1-256) and the his-EmGFP-DBNL(1-374) slowed down significantly at 10 nM compared to 100 pM concentration (Fig. 3F and G). Thus, it can be inferred that at 10 nM concentration isolated ADF-H domain exists as a monomer, while both mutants truncated after coiled-coil domain undergo oligomerization at the same concentration. These results indicate that the coiled-coil domain mediates self-assembly, but the presence of both proline-rich and SH3 domains is essential for DBNL to maintain stable dimer conformation. It is tempting to speculate that oligimerization of ADF-H domain of DBNL is concentration-dependent, similar to that of human cofilin, which consists of a single ADF-H domain and displays a concentration-dependent self-assembly via its C-terminal part^32,33^.

### *In vivo* interaction of DBNL and truncated DBNL mutants with *β*-actin probed with FLIM-FRET

Previously, we used atomic force microscopy to show that drebrin-like protein has a globular shape and decorates the sides of actin filaments^3^. Using an *in vitro* actin-binding assay, Kessels and colleagues identified two independent actin-binding modules within the structure of the mammalian DBNL: ADF-H and coiled-coil domains^1^. In a two-component *in vitro* system, DBNL truncated after a coiled-coil domain binds to F-actin as strong as the full-length protein, while the isolated ADF-H domain binds with significantly reduced affinity^1^. To probe the impact of protein conformation on its interaction with F-actin inside cells, we utilized FLIM-FRET imaging^34–36^. In FLIM-FRET, one labels the target molecule of interest with a donor, and the corresponding ligand with an acceptor, and then determines the FRET efficiency between donor and acceptor by measuring the donor fluorescence lifetime. Due to the short nanometer range of FRET, a high FRET efficiency indicates direct target-ligand binding. A reasonable control experiment is to measure the fluorescence lifetime of donor in the absence of acceptor, or by re-measuring the donor lifetime after full acceptor photobleaching. For our purpose, we co-expressed EGFP (enhanced GFP)-actin (donor) and mCherry-DBNL or mCherry-tagged truncated DBNL mutants (acceptor) in MCF7 cells and measured the fluorescence lifetime of EGFP-actin.

FLIM-FRET experiments show that the average lifetime of EGFP-actin inside cells in presence of mCherry-DBNL is shorter compared to cells expressing only EGFP-actin (donor only). This finding indicates a strong interaction of both proteins (Fig. 4A and E). In contrast, average lifetime values of EGFP-actin in cells that additionally express mCherry-tagged DBNL(1-179), DBNL(1-276) or DBNL(1-374) (Fig. 4B-D) match closely ‘donor only’ cells (Fig. 4E). The absence of a FRET-induced lifetime reduction of EGFP-actin in the presence of DBNL mutants suggest that they do not bind to actin as the full-length DBNL. Alternatively, other actin-binding proteins with higher F-actin-binding affinity might displace truncated DBNL from actin filaments in live cells. In any case, the experimentally observed functional properties of truncated DBNL clearly indicate the importance of its native structure for unaltered biological activity. In cells, DBNL undergoes proteolysis by the ubiquitous calcium-sensitive protease calpain-2, which cleaves DBNL between the coiled-coil and the proline-rich region^5^. Here, the N-terminal fragment (consisting of two actin-binding modules) alone is not able to rescue formation of actin-based dorsal ruffles in DBNL-deficient cells^5^. The expression of ADF-H domain alone is known to increase Rho GTPase signaling and to induce breast cancer cell invasion^8^. In *C. elegans*, the impact of truncations can be visually evaluated by the change in worm’s movement ability. While the *dbn-1(vit7)* mutant, containing a premature stop codon after the largest part of ADF-H domain, displays a strong body curvature and a striking hyper-bending phenotype, the truncated after two coiled-coils *dbn-1(ok925)* mutant moves in sinusoidal waves similar to the wild type (submitted, Butkevich et al.). However, the *ok925* has a mild disorganization of actin filaments during body-wall muscle contraction^3^ and defective vesicle scission during endocytosis in the intestines^37^. Taken together, our data demonstrate that the amino-acid sequence of DBNL defines its existence as a dimer and that DBNL can efficiently interact with actin filaments inside a cell only in its native conformation.

**Figure 4:**
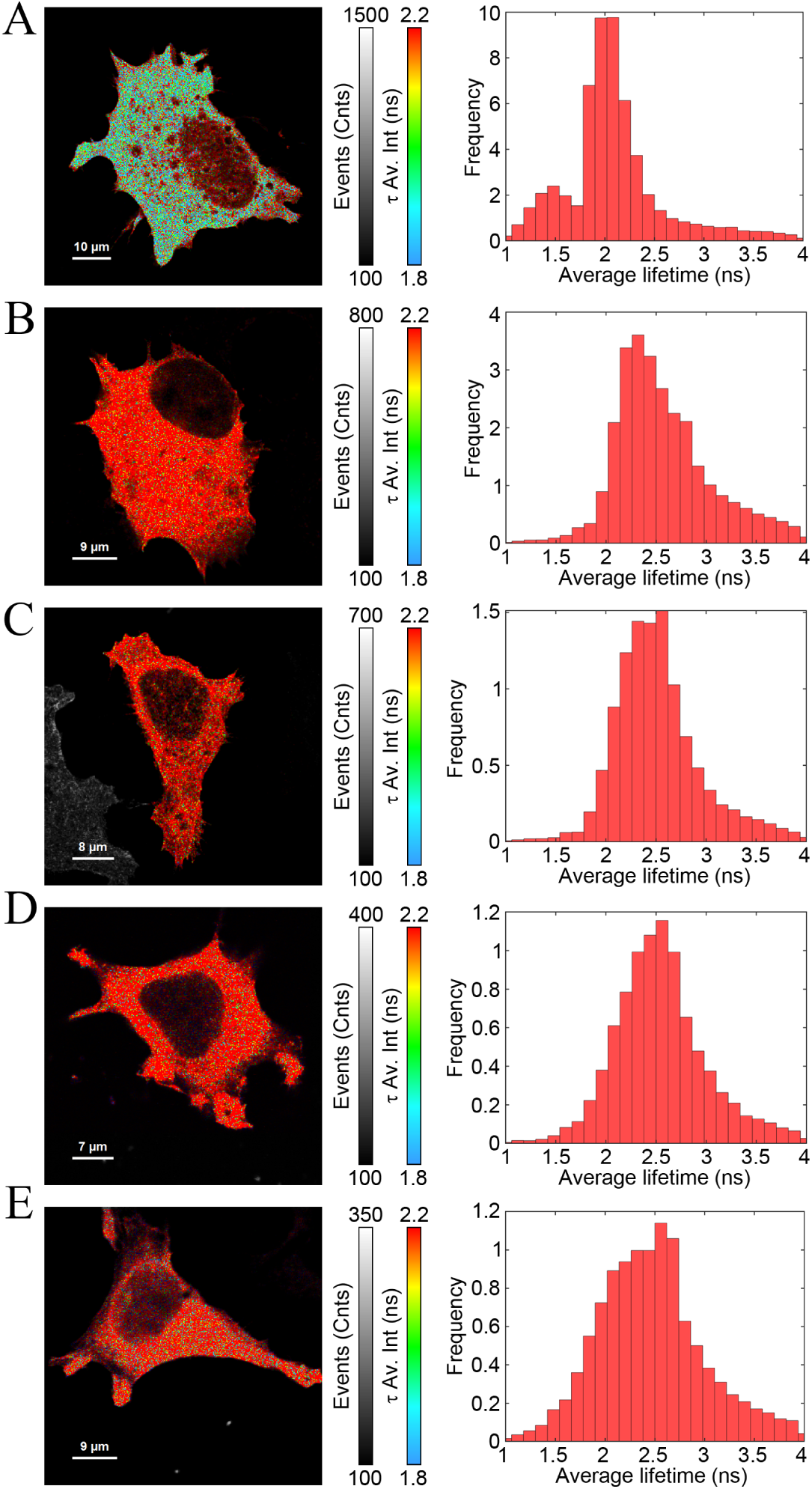
Interaction of EGFP-actin and mCherry-DBNL and its truncation mutants measured using FLIM-FRET imaging. **A**. MCF7 cells were co-transfected with EGFP-actin (donor) and mCherry-DBNL (acceptor). Fluorescence intensity weighted average lifetime image (left panel) and the corresponding lifetime histogram of the donor are shown (right panel). **B**. The same for EGFP-actin and mCherry-DBNL(1-179). **C**. The same for EGFP-actin and mCherry-DBNL(1-256). **D**. The same for EGFP-actin and mCherry-DBNL (1-374). **E**. The same for ‘donor only’ sample – EGFP-actin alone. As can be seen the average fluorescence lifetime of EGFP-actin in presence of mCherry-DBNL (**A**) is shorter (1.5-2 ns) compared to the ‘donor only’ (**E**) (2.3-2.5 ns) and the truncated mutants samples (**B**-**D**). Colorbar legend ‘*τ* Av.int (ns)’ and ‘events (counts)’ represent intensity weighted average lifetime and intensity in photon counts respectively.

## Materials and Methods

### Expression constructs

A cDNA encoding human DBNL was amplified by PCR from Human Brain Whole QUICK-Clone™ cDNA (Clontech Laboratories) using the primers: attB1DBNL forward 5’-GGGGAC AAGTTTGTACAAAAAAGCAGGC TCGATGGCGGCGAACCTGAGCCG-3’ and attB2DBNL reverse 5’–GGGGACCACTTTGTACAAGAAAGCTGGGTGTCACTCAATGAGCTCCACGT-3’, subcloned into pDONR201 (Invitrogen) and subsequently cloned into CMV-mCherry-Gateway vector or Vivid Colors™ pcDNA™6.2/N-EmGFP-DEST vector (ThermoFisher Scientific) to obtain CMV-mCherry-DBNL or CMV-EmGFP-DBNL, respectively. To create a CMV-mCherry-Gateway vector, mCherry was amplified from pmCherry-N1 vector (Clontech Laboratories) and placed between XhoI and XbaI sites to exchange EmGFP in pcDNA™6.2/N-EmGFP-DEST. cDNAs encoding DBNL, mCherry-DBNL, EmGFP-DBNL, mCherry and EmGFP alone as well as EmGFP-tagged DBNL(1-179), DBNL(1-256) and DBNL(1-374) were amplified by PCR and cloned into pET28a vector (Novagen) between BamHI and NotI sites to obtain constructs for expression of his-tagged proteins in *E. coli*. CMV-Gateway vector was created by excision of EmGFP from pcDNA™6.2/N-EmGFP-DEST between two XhoI sites. cDNAs encoding DBNL, DBNL(1-179), DBNL(1-256) and DBNL(1-374) were amplified by PCR and cloned into CMV-Gateway vector and into CMV-mCherry-Gateway vector for expression in MCF7 cells. All cDNA clones were confirmed by DNA sequencing at SeqLab. pEGFP-Actin (6116-1) was purchased from Clontech Laboratories.

### Expression and purification of recombinant proteins

*E. coli* BL21-Gold (Agilent Technologies) were transformed with pET28a-mCherry-DBNL, pET28a-EmGFP-DBNL, pET28a-EmGFP-DBNL(1-179), pET28a-EmGFP-DBNL(1-256), pET28a-EmGFP-DBNL(1-374), pET28a-EmGFP, or pET28a-mCherry. Cells were grown to an OD_600_ of 0.4 in in Luria-Bertani medium containing 50 mg/l kanamycin at 37 °C. Cells were induced to express protein by the addition of 0.1 mM IPTG, and incubation continued for a further 4-5 h at 22 °C. Cultures were centrifuged at 4,600 g for 20 min at 4 °C. Pellets were suspended in 20 ml of lysis buffer containing 20 mM NaH_2_PO_4_ pH 8.0, 500 mM NaCl, 10 mM *β*-mercaptoethanol, 1 mg/ml lysozyme, 0.01 mg/ml DNAse and protease inhibitor cocktail (S8830, Sigma-Aldrich) and subjected to three times alternating procedures of freezing in liquid nitrogen followed by thawing on ice. Samples were sonicated 4 times 20 second each on melting ice with intervals for cooling using a Branson model 250 sonifier. Lysates were spun for a further 20 min at 40,000 g at 4 °C and resulting supernatants were incubated 1 hour with Ni^2+^-nitrilotriacetate (NTA) agarose beads (QIAGEN) at 4 °C on rotating wheel. Beads were washed in buffer containing 20 mM NaH_2_PO_4_ pH 8.0, 500 mM NaCl, 10 mM *β*-mercaptoethanol, 40 mM imidazole, 1mM PMSF. Protein was eluted with buffer containing 20 mM NaH_2_PO_4_ pH 8.0, 500 mM NaCl, 10 mM *β*-mercaptoethanol, 350 mM imidazole, 1mM PMSF. Elution buffer was exchanged to PBS pH 7.4 (D8537, Sigma-Aldrich) using Amicon Centrifugal Filters (Merck).

### Antibodies

Antibodies specific for DBNL were raised in chicken against his-tagged full-length human DBNL coupled to keyhole limpet hemocyanin at Bioscience. The final antisera was purified using NHS-activated Sepharose High Performance (GE Healthcare Life Science). The reason for generation of these antibodies was that two tested anti-DBNL antibodies (sc398351, Santa Cruz and HPA020265, Sigma-Aldrich) recognized a denatured but not a native form of DBNL and any antibodies, which recognize native human DBNL were reported so far. The antibodies were validated in western blot. The immunoreactivity was significantly reduced upon treatment of MCF7 cells with shRNA against human DBNL (sc-75255-SH, Santa Cruz Biotechnology). The band was not detected by this antibodies preincubated with his-DBNL and by preimmune serum subjected to purification in parallel to postimmune serum. As seen in Fig. 3B, these antibodies recognize full-length DBNL and its truncation mutants expressed from the plasmids. The signal from the truncated construct DBNL(1-179) is reduced presumably due to the reduction of number of antibody-binding sites.

### Cell culture and transfection

MCF7, HEK-293 and HeLa cells were purchased from ATCC and hMSC from Lonza. All cell lines tested negative for micoplasma contamination. Cells were cultured in DMEM supplemented with 10% FCS, 2 mM L-glutamine, 1 mM sodium pyruvate, 100 U/ml penicillin-streptomycin in a humidified 5% CO_2_ atmosphere at 37 °C. For western blot, MCF7 cells were transfected with non-tagged DBNL, DBNL(1-179), DBNL(1-256) or DBNL(1-374) using ViaFect (Promega) and assayed 24 h later. For FLIM-FRET imaging, MCF7 cells were transfected with EGFP-actin alone or co-transfected with EGFP-actin and mCherry-DBNL, mCherry-DBNL(1-179), mCherry-DBNL(1-256) or mCherry-DBNL(1-374). 12 hours later cells were rinsed with PBS and fixed in 4 % (v/v) formaldehyde/PBS for 15 min. Samples were washed in PBS, rinsed with distilled water, and mounted using Fluoroshield™ (F6182, Sigma-Aldrich).

### Gel electrophoresis and western blot

About 200,000 cells were lysed in 100 *µ*l of lysis buffer, containing 150 mM NaCl, 50 mM Tris pH 7.4, 2 mM EDTA and 1% Nonidet P-40. For native gel electrophoresis, 2x Tris-Glycin buffer was added and 20 *µ*l of sample were resolved by 10% Novex Tris-Glycin gel (Life Technologies). For SDS PAGE, 4x Laemmli buffer was added, the sample was heated to 95 °C for 5 min and 15 *µ*l of sample were resolved by a 12% NuPAGE Bis-Tris gel (Invitrogen). Proteins were transferred onto a Immobilon-P polyvinylidene difluoride membrane (Millipore) using a Bio-Rad Criterion Blotter. After the transfer membrane was blocked with 10% non-fat milk in PBS containing 0.05% Tween 20 for 30 min and incubated with affinity-purified anti-DBNL antibodies diluted at 1:1000 at 4 °C overnight, followed by treatment with diluted at 1:10000 peroxidase-labeled goat anti-chicken IgY (A16054, Thermo Fisher). Mouse monoclonal anti-actin (clone C4; MP Biomedicals) and mouse monoclonal anti-*α*-tubulin (clone DM1A, Sigma-Aldrich) antibodies were used in combination with peroxidase-labeled goat anti-mouse IgG (A4416, Sigma-Aldrich). The reactivity was detected with an Amersham ECL Detection Reagent (GE Healthcare Life Science) using an Intas gel imager (Intas Science Imaging Instruments GmbH). To analyze the electrophoretic mobility of recombinant proteins, 2 *µ*M solution of each protein in native sample buffer or in Laemmli buffer were prepared. For native gel electrophoresis, 15 *µ*l of the sample were resolved by 10% Novex Tris-Glycin gel (Life Technologies). For SDS PAGE, the sample was heated to 95 °C for 5 min and 15 *µ*l of the sample were resolved by 12% NuPAGE Bis-Tris gel (Invitrogen). Molecular weight markers were purchased from ThermoFisher Scientific (PageRuler™ Plus Prestained Protein Ladder) and Serva (Native Marker 39219). For detection of native markers, a PVDF membrane was stained with Ponceau S (Sigma-Aldrich). Gels were stained with Roti^®^-Blue quick (Roth).

### Instrumentation and experimental procedures for FCCS, 2fFCS, FCS and FLIM-FRET imaging

For FCS, 2fFCS and FCCS experiments, we used the commercial instrument Microtime 200 (PicoQuant GmbH)^38^.

#### FCS

Excitation was done with a linearly polarized pulsed diode laser (*λ* = 485 nm, pulse duration 50 ps FWHM, LDH-P-C-485B, PicoQuant) equipped with a clean-up filter (Brightline FF01-480/17, Semrock). Light of this laser is pulsed at a repetition rate of 40 MHz with a multi-channel picosecond laser driver (PDL 828, “Sepia II”, PicoQuant). The laser beam is coupled into a polarization-maintaining single-mode fiber (PMC-400-4.2-NA010-3-APC-250V, Schäfter and Kirchhoff GmbH). At the fiber output, the light is collimated and reflected by a dichroic mirror (FITC/TRITC Chroma Technology) into the objective lens of the microscope (UPLSAPO 60x water, 1.2 N.A., Olympus). The same water-immersion objective is used to collect fluorescence from the sample. A long-pass filter (BLP01-488R-25, Semrock) is used to block back-scattered light from the laser. The emission light is focused into a pinhole of 100 *µ*m diameter, collimated again, and split by a non-polarizing beam splitter cube (Linos Photonics GmbH & Co. KG) and refocused onto two single-photon avalanche diodes (SPCM-CD 3516 H, Excelitas Technologies GmbH & Co. KG). A multichannel picosecond event timer (HydraHarp 400, PicoQuant) records the detected photons from both detectors independently with an absolute temporal resolution of 16 ps on a common time frame. For FCS measurements, his-EmGFP-DBNL(1-179), his-EmGFP-DBNL(1-256) and his-EmGFP-DBNL(1-374) were diluted to 10 nM and 100 pM concentration in PBS (D8537, Sigma-Aldrich). The experiments were done by putting 30 *µ*l of sample on top of a cleaned glass cover-slip, and during measurements the objective was focused approximately 30 *µ*m into the solution.

#### FCCS

The excitation unit consists of two pulsed diode lasers, blue (*λ*= 485 nm, LDH-P-C-485B, PicoQuant) and red (*λ*exc = 640 nm, LDH-D-C 640, PicoQuant), for his-EmGFP-DBNL and his-mCherry-DBNL, respectively. Laser pulse width was 50 ps FWHM, and pulse repetition rate was 40 MHz. Laser pulsing, beam coupling, and focusing through the objective lens were done as described in the previous section. After the pinhole, the collected emission light was split by a non-polarizing beam splitter cube (Linos Photonics GmbH & Co. KG) and refocused onto two single-photon avalanche diodes (SPCM-CD 3516 H, Excelitas Technologies GmbH & Co. KG). Spectral cross-talk between green and red channel was avoided using emission band-pass filter 525/45 (Semrock) and 692/40 (Semrock) in front of each detector. His-EmGFP-DBNL and his-mCherry-DBNL were diluted in PBS (D8537, Sigma-Aldrich) to 1 nM each and 30 *µ*l of mixed sample solution was placed on top of a cleaned glass cover-slip for measurement. The same was done for his-EmGFP and his-mCherry. Excitation and detection for 2fFCS were done as described in^22^.

#### FLIM-FRET Imaging

For FLIM-FRET experiments, we used a home-built confocal setup equipped with an objective lens of high numerical aperture (Apo N, 100× oil, 1.49 NA, Olympus Europe, Hamburg, Germany). The excitation unit consists of a pulsed, linearly polarized white light laser (SC400-4-80, Fianium Ltd., pulse width ∼ 50 ps, repetition rate 80 MHz) equipped with a tunable filter (AOTFnC 400.650-TN, Pegasus Optik GmbH). For our experiments, we used 488 nm wavelength for exciting EGFP tagged to actin filaments. Light was reflected using a non-polarizing beam splitter towards the objective lens, and the back-scattered excitation light was blocked with long-pass filters (EdgeBasic BLP01-488R, Semrock). A band-pass filter 525/45 (BrightLine FF01-525/45, Semrock) was used for our measurements. Emission light collected by the objective was focused into a pinhole of 100 *µ*m diameter, collimated again and refocused into the active area of an avalanche photodiode (Excelitas Technologies Corporation). Data recording was performed with the aid of a multichannel picosecond event timer (HydraHarp 400, PicoQuant GmbH). Individual cells were scanned with a focused laser spot using a piezo-nanopositioning stage (P-562.3CD, Physik Instrumente GmbH).

### Data evaluation

Calculation of intensity autocorrelation and cross-correlation curves for FCCS were done as described in^16,17^ using a custom written MATLAB routine. For FCS, fluorescence correlation curves were calculated from intensity fluctuations using the algorithm of^39^. Finally, data evaluation and fitting for 2fFCS were performed as described in^22^ utilizing again custom written MATLAB routines. Fluorescence lifetime data evaluation was done using MATLAB software package for lifetime fitting as available here – https://www.joerg-enderlein.de/software.

## Acknowledgements

We are grateful to Prof. Dr. Christoph F. Schmidt for providing lab facilities and resources for protein expression and purification. We thank Tanja Gall for technical assistance. Funding was provided by the Deutsche Forschungsgemeinschaft (DFG) via project A06 of the Collaborative Research Center SFB 860 and the European Research Council under the European Union’s Seventh Framework Programme (FP7/2007-2013) / ERC grant agreement n°340528.

## Author contributions

AG did the FCS, FCCS, 2fFCS, and FLIM-FRET experiments, data analysis and co-wrote the manuscript. JE co-wrote the manuscript and helped in data analysis. EB designed the project, generated the constructs, performed cell culture and biochemical experiments, and co-wrote the manuscript.

